# *Escherichia coli* adaptation under prolonged resource exhaustion is characterized by extreme parallelism and frequent historical contingency

**DOI:** 10.1101/2024.03.21.586114

**Authors:** Shira Zion, Sophia Katz, Ruth Hershberg

## Abstract

Like many other non-sporulating bacterial species, *Escherichia coli* is able to survive prolonged periods of resource exhaustion, by entering a state of growth called long-term stationary phase (LTSP). In July 2015, we initiated a set of evolutionary experiments aimed at characterizing the dynamics of *E. coli* adaptation under LTSP. In these experiments populations of *E. coli* were allowed to initially grow on fresh rich media, but where not provided with any new external growth resources since their establishment. Utilizing whole genome sequencing data obtained for hundreds of clones sampled at 12 time points spanning the first six years of these experiments, we reveal several novel aspects of the dynamics of adaptation. First, we show that *E. coli* continuously adapts genetically, up to six years under resource exhaustion, through the highly convergent accumulation of mutations. We further show that upon entry into LTSP, long-lasting lineages are established. This lineage structure is in itself convergent, with similar lineages arising across independently evolving populations. The high parallelism with which adaptations occur under LTSP, combined with the LTSP populations’ lineage structure, enable us to screen for pairs of loci displaying a significant association in the occurrence of mutations, suggestive of a historical contingency. We find that such associations are highly frequent and that a third of convergently mutated loci are involved in at least one such association. Combined our results demonstrate that LTSP adaptation is characterized by remarkably high parallelism and frequent historical contingency.

**Author summary:** Characterizing the dynamics by which adaptation occurs is a major aim of evolutionary biology. Here, we study these dynamics in five populations of *Escherichia coli*, independently evolving over six years under conditions of resource exhaustion. We show that even under very prolonged resource exhaustion bacteria continuously genetically adapt. Within our populations long lasting lineages are established, each of which undergoes independent and continuous adaptation. We demonstrate that bacterial adaptation under resource exhaustion is both highly convergent – meaning that same adaptive mutations occur across independently evolving populations and lineages, and frequently historically contingent – meaning that the adaptive nature of many of the adaptations we see depends on the previous occurrence of other adaptations.

## Introduction

Evolution is a historical process, in which the potential of certain adaptive events to occur, may sometimes depend on the previous occurrence of other events. Historical contingency is generally defined as the dependence of adaptation on the history of a population [1].

Such history can manifest through the chronology of experienced conditions, the order of previously acquired mutations, or both [2–4]. The role of historical contingency in determining adaptive trajectories has long been debated. Researchers have argued about whether evolution is largely deterministic, most frequently following the same paths, or whether evolution tends to be far less predictable, due in part to frequent contingencies, making it impossible to “replay life’s tape” [5,6]. Making these arguments more confusing, the word ‘contingency’ itself is often used to imply different meanings: on the one hand a possible, but quite unlikely and thus unpredictable, future event (e.g. planning for every contingency), and on the other hand an event whose occurrence depends on a different event and is quite likely given that event. In this paper, the term ‘contingency’ will be used to describe mutations whose adaptive nature is significantly dependent on the previous acquisition of other adaptive mutations.

One well characterized example of historical contingency was revealed as part of Richard Lenski’s seminal long-term evolutionary experiment (LTEE) [7,8]. The LTEE was initiated in 1988, when 12 independent populations of *E. coli* were established in glucose-limited growth media, supplemented with citrate. Since 1988, these populations have been daily diluted into fresh media, allowing for the characterization of the dynamics of adaptation of *E. coli* under the experiment’s conditions of limited glucose, for well over 80 thousand generations. *E. coli* is unable to utilize citrate for carbon under oxic conditions [8,9]. Indeed, this inability is one of the key parameters used to distinguish *E. coli* from other bacterial species [10]. One, and only one of the 12 LTEE populations evolved the ability to consume citrate as a carbon source, following ∼30,000 generations. Further analyses revealed that the reason that this phenotype evolved in only a single population, following a fairly long period of time, was related to historical contingency. Specifically, the ability of *E. coli* to adapt to utilize citrate as a carbon source required a large number of mutations to occur in a fairly specific order, with the effect of certain mutations depending on the previous occurrence of others [8,11–14].

Fitting with the instance of contingency identified in the LTEE, historical contingency is often framed as a factor that reduces the predictability of adaptive trajectories [3,15,16]. After all, if specific mutations are adaptive only in individuals carrying other specific adaptations, and if many different adaptations can initially occur, different lineages will likely adapt via divergent trajectories. Furthermore, such trajectories may become more and more divergent as additional adaptations accumulate, further reducing the ability to predict how a given population will adapt. At the same time, many evolutionary experiments reveal a strong tendency for adaptation to occur in a convergent manner [17–19]. Such convergence, or parallelism (the terms will be used interchangeably in this paper), means that similar adaptive patterns and events tend to happen in parallel, across independently evolving populations, implying increased predictability. It has been shown for example that independently evolving LTEE populations displayed similar trajectories when it came to the rates with which they improved their fitness (a pattern of diminishing returns with time) [20,21].

Additionally, different LTEE populations often experienced similar phenotypical changes [7,8,20,22,23]. Finally, convergent mutation acquisition was also observed in the LTEE, with different populations often accumulating mutations to similar loci [8,22,24–26].

In July of 2015, we initiated our own long-term evolutionary experiment, designed to study *E. coli* adaptation under prolonged resource exhaustion [27,28]. *E. coli* and many other non-sporulating bacteria are able to survive long periods of resource exhaustion, following brief periods of growth, by entering a state of growth termed long term stationary phase (LTSP) [27,29–32]. In our LTSP experiment, five populations of *E. coli* K12 MG1655 were established by inoculating cells into 400 ml of fresh rich media. These populations have not been provided with any external growth resources ever since and have had to survive by recycling the remains of their ancestors and brethren [27,33]. After a day of growth these populations entered a period of rapid death, followed by entry into LTSP, at about day 11 of the experiment. Since day 11 of our experiments, we have seen a slow decline in viability with time, and populations remain viable to this day.

Data from the sequencing of ∼10 clones per population, from each of nine sampled time points, spanning the first three years of the experiment (for a total of 413 sequenced clones), have enabled us to begin characterizing the dynamics of adaptation under LTSP [28]. These analyses have revealed that during the studied timeframe, LTSP populations continuously and rapidly adapt through the continuous accumulation of mutations. Such adaptation occurs in a highly convergent manner, with mutations to the same loci and often even to the exact same sites very frequently occurring across independently evolving populations. In three of the five populations we saw the emergence of mutators, that having acquired a mutation to a mismatch repair gene, substantially increased their mutation rates. Such mutators coexisted within the same populations, alongside non-mutators, for extended periods of time [28].

Here we expand on our previously available data by sequencing clones sampled up to six years into our experiment. Utilizing these data, we characterize the lineage structure of each of our five LTSP populations. We show that the lineage structure of LTSP populations is itself a convergent feature of LTSP adaptation. This lineage structure, together with the extreme convergence of adaptation under LTSP enables us to search for putative instances of historical contingency (**Fig 1**). Results of this analysis reveal that historical contingencies appear to be very common, as we find that more than a third of convergently mutated loci are involved in a putative contingency with at least one additional convergently mutated locus.

**Fig 1.**
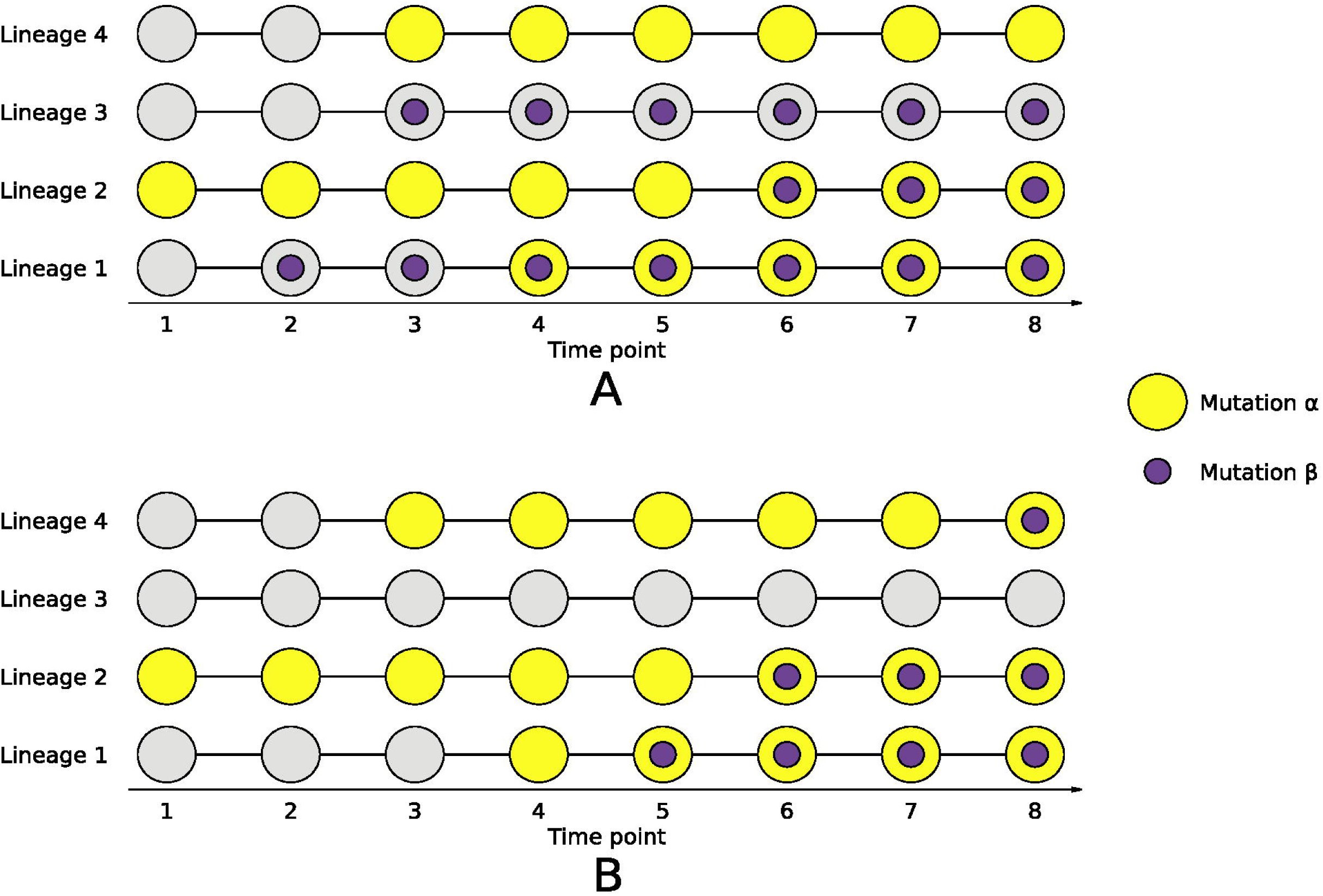
Schematic representation of how patterns of occurrence of convergent mutations can enable the identification of putatively contingencies. Two options are schematically presented: (A) An independent mutation acquisition pattern does not suggest a contingency relationship. (B) Strong association between the occurrence of mutations suggests a putative contingency. In the presented example it would be possible to predict that mutation is contingent on the previous occurrence of mutation .

## Results

### Continuous convergent adaptation under LTSP

Our five LTSP populations, established July 2015 maintain their viability to this day. **Fig 2A** depicts viability as function of time, as estimated by plating samples and quantifying colony forming units (CFUs). We previously sequenced 413 clones from 9 time points spanning the first three years of our LTSP experiments. Here, we add to these data, resulting in whole genome sequencing data of 637 clones sampled at 12 time points, spanning the first 6 years of the experiment. A full list of the sequenced clones and their obtained sequencing coverage is given in **S1 Table**. A full list of called mutations is provided in **S2 Table**. Focusing on 392 non-mutator clones, **Fig 2B** depicts the number of mutations accumulated as a function of time spent under LTSP. As can be seen, mutations continue to accumulate with time, throughout the six years examined so far.

**Fig 2.**
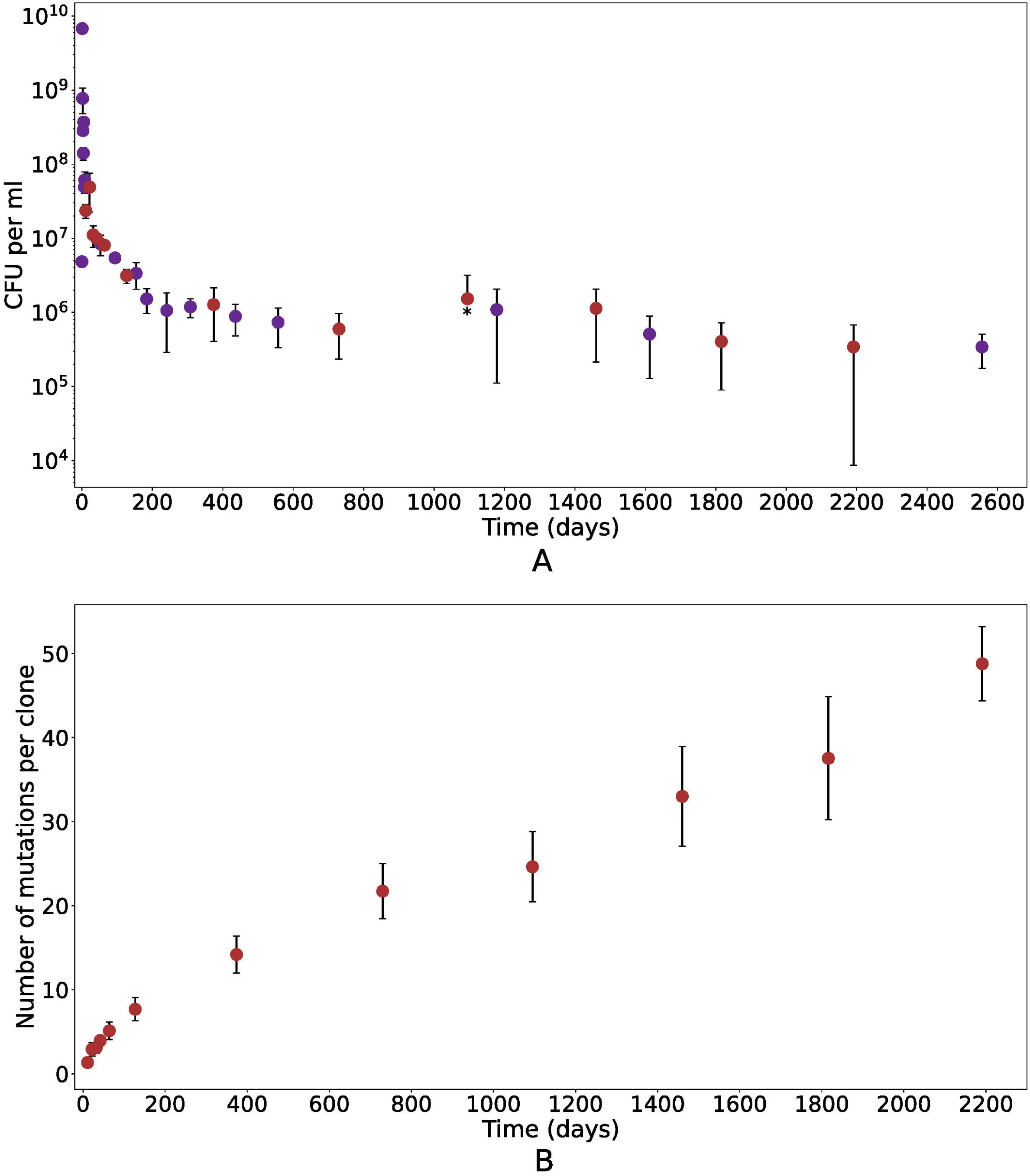
Viable cell counts and mutation accumulation in five *E. coli* LTSP populations. **(A)** mean colony forming unit (CFU) counts across the five independent populations of *E. coli*, during the first seven years under LTSP. The 12 time points at which ∼10 clones per population were fully sequenced are marked by red dots. **(B)** The mean number of mutations acquired by non-mutator clones, as a function of time. For both plots, error bars represent standard deviations around the presented means. An asterisk marks a negative standard deviation that could not be plotted on a logarithmic scale since it was larger than the mean.

Mutations accumulated under LTSP within non-mutator clones are significantly enriched for non-synonymous vs. synonymous mutations, relative neutral expectations (dN/dS = 3.5, *P* << 0.001, according to a χ^2^ test). This indicates that mutation accumulation within non-mutators is governed by positive selection [26,34,35]. An additional indication that many of the observed mutations are adaptive is the extremely high convergence with which mutations within non-mutator clones occur. ∼88.5% of mutations acquired by non-mutator clones, fall within 265 loci that are mutated in at least two independently evolving populations (**S1A Fig**, **S3 Table**). These 265 loci constitute only ∼5.7% of the *E. coli* genome. Combined, these results show that our LTSP populations continuously adapt in an extremely convergent manner, up to at least six years under resource exhaustion.

### Lineage structure and convergent lineage dynamics under LTSP

One of the most striking examples of convergence that we already reported on in our previous publications involves mutations in the RNA-polymerase core enzyme (RNAPC)-encoding genes *rpoB* and *rpoC* [27,28]. Strikingly, more than 90% of all sequenced clones carry such a mutation, with the same specific mutations often appearing in a convergent manner across populations, and with clones carrying different RNAPC adaptations often co-existing within single populations (**Fig 3A**). The RNAPC mutations are often the first mutations to occur, and thus, all of the descendants of the clones that initially acquired these specific *rpoB* and *rpoC* mutations also carry them, and most other mutations are acquired on the background of these RNAPC mutations. It can thus be said that the occurrence of the RNAPC mutations early on in LTSP establishes lineages, that then co-exist alongside other lineages, established from clones carrying different RNAPC lineage-defining mutations.

**Fig 3.**
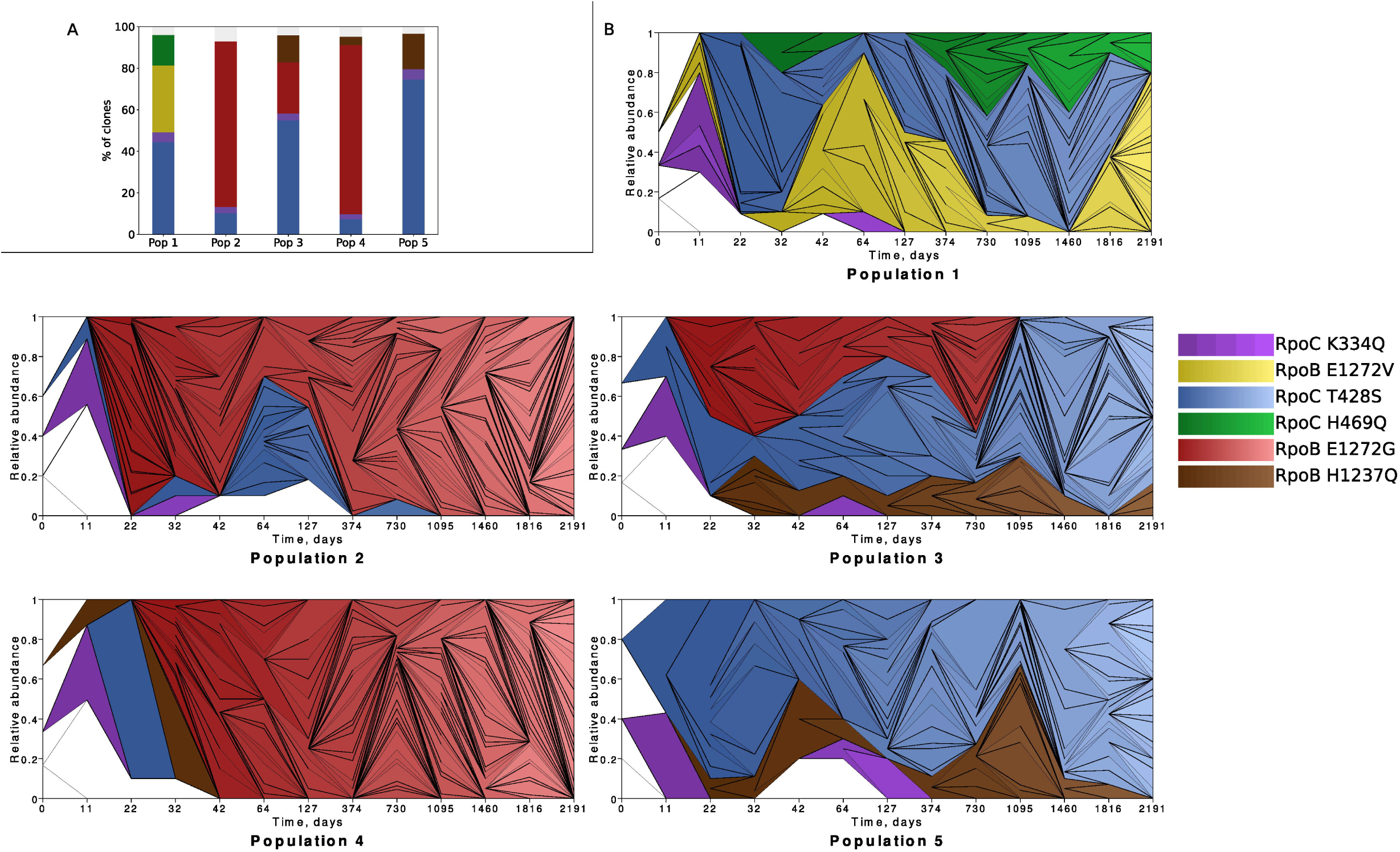
Convergent lineage structure under LTSP. **(A)** Convergent pattern of RNAPC lineage-defining mutation acquisition across populations. **(B)** Muller plots depicting the relative abundance of the different lineages, as a function of time. Lineages are colored according to their lineage defining mutations, allowing for the use of the same colors in both panels A and B.

By using our WGS data to build phylogenies within each population we characterized each population’s lineage structure (see **Materials and Methods**). Obtained results were visualized as Muller plots depicting the relative frequencies of lineages as a function of time (**Fig 3B**). These Muller plots were drawn with the help of the R package MullerPlot [36]. To build our phylogenies and characterize our lineages we focused on mutations that reached at least 30% frequency during at least one timepoint. For a lineage to be defined, it was required that the phylogeny of a potential lineage lasted within our population for at least two time points, and had been founded by a mutation that, at some timepoint reached at least 30% frequency within its population. We were able to assign ∼95.3% of all sequenced clones to a lineage. A total of 18 lineages were identified across our five populations. While this was not an a-priori requirement, for all lineages the founding mutation was one of the RNAPC mutations summarized in **Fig 3A**. We observed continuous mutation accumulation occurring with time within lineages (**S2 Fig**). This means that clones from a specific lineage sampled at a given time point tended to accumulate additional mutations, compared to clones sampled from the same lineage at earlier time points.

Interestingly, lineages that share the same lineage-defining mutation, but belong to different populations often behave similarly. The most striking example of this involves lineages defined by the RpoC K334Q mutation, which emerge early on across all five populations, but then also reduce in frequency below our detection ability by day 127, across all populations (**Fig 3B**). In contrast, lineages defined by RpoC T428S also establish early in all five independent populations, but persist at high frequencies over the duration of the experiment so far, in all three populations that were not taken over by mutators.

Mutations within the same populations occurring within clones belonging to different lineages have necessarily occurred independently. We were therefore able to examine convergence not only between populations, but also within them. Our five populations contain 18 independently evolving lineages. ∼89.3% of non-mutator mutations fall within 268 loci that are mutated in two or more of these lineages (**S1B Fig**, **S4 Table**). Other than the RNAPC genes *rpoB* and *rpoC*, the most convergently mutated locus is the gene encoding the global transcription regulator CRP, which is mutated in 11 of 18 independently evolving lineages. Mutations occurring within genes mutated in two or more lineages are significantly enriched for non-synonymous vs. synonymous mutations, relative random expectations (dN/dS = 5.5, *P* << 0.001, according to a χ^2^ test), while, in contrast mutations within loci that are mutated in only a single lineage are enriched for synonymous mutations (dN/dS = 0.4, *P* << 0.001, according to a χ^2^ test). This suggests that loci mutated in two or more lineages are accumulating LTSP mutations as a result of positive selection [26,34,35]. Such convergently mutated loci are significantly enriched for genes involved in regulation of gene expression and transport (see **Materials and Methods**).

### Frequent association between the occurrence of mutations across loci

Once we determined the lineage structure of our populations and assigned lineages according to the RNAPC mutation that established them, it became evident that mutations within certain loci occurred predominantly within lineages established by a specific RNAPC mutation. The most striking example of this is that mutator clones always belong to the lineages defined by the RpoB E1272G mutation. The RpoB E1272G mutation established lineages in three of our five populations, and across all these populations, all clones sequenced belonging to these lineages also had a mutator mutation, within a mismatch repair gene (**S2 Table**). In two of the populations (populations #2 and #3) mutators appear to have emerged only once, as all mutator clones within these populations carried the exact same mismatch repair gene mutation. In the third population (population #4) two different types of mutators evolved. However, both evolved on the background of population 4’s RpoB E1272G lineage. No other lineages ever included any such mutator clones. Interestingly, the pattern by which mutators occur only on the background of lineages established by the RpoB E1272G mutation was also recapitulated in a second study, in which six additional LTSP populations were established [37]. Two additional examples of loci that are mutated predominantly within lineages defined by a specific RNAPC mutation are: (1) the emergence of mutations in both the *dctA* dicarboxylate transport gene and its promoter on the background of three out of five lineages defined by the RpoC K334Q mutation, and never on the background of any other lineage-defining mutations; and (2) the emergence of specific mutations within the gene *dppA* and the *gcvT* promoter as well as different nonsense and frameshift mutations to *proQ* that occur in four of five lineages defined by the RpoC T428S mutation (the fifth RpoC T428S lineage that did not acquire these mutations was very short-lived, due to the RpoB E1272G-muator lineage overtaking its population by day 42 of the experiment).

Next, focusing on all 268 loci that are mutated in at least two independently evolving lineages or populations (excluding *rpoB* and *rpoC*), we extracted all possible loci pairs for which there existed clones that carried mutations in both loci in at least two independently evolving lineages or populations (see **Materials and Methods**). For each such pair of loci, we then asked in how many lineages did we see clones that carried mutations to both loci, clones that carried mutations to neither locus, and clones that carried mutations in only one or only the other of the two loci. Using these counts, we could then calculate the strength of association between the presence of mutations in one locus and the presence of mutations in the other, across all lineages. We calculated three coefficients of association which range in their strictness from low to high: Yule’s association coefficient Q [38], Yule’s coefficient of colligation Y and Yule’s phi coefficient of contingency φ [39]. Depending on the coefficient used we identified between 360 and 664 instances of statistically significant association (after correcting for multiple testing, see **Materials and methods**, **S5 Table**). In total, we found that 91-94 out of the 268 (34.0-35.1%) convergently mutated loci are involved in at least one significantly associated loci pair. It can thus be said that for more than a third of convergently mutated loci the presence of mutations is significantly associated with the presence of mutations within at least one additional locus. To further support the biological relevance of the predicted contingent loci pairs, we examined whether they tended to be functionally related, more often than expected by chance, according to annotations contained within the STRING database [40]. Depending on the coefficient of association used, between 18.1% and 21.7% of putatively contingent loci pairs had known functional relationships. Based on 1000 repeated randomizations of our networks of putatively contingent loci pairs, we found that for all three coefficients, this proportion was significantly larger than expected by chance (see **Materials and Methods**, **S6 Table**).

The different coefficients used vary greatly in their stringency. As a result, many more putative contingent pairs were identified for the less strict coefficients. We aimed to set cutoffs for each of the coefficients that would allow us to select the most reliable putative contingencies, without discarding too many meaningful pairs. To do so, for each of the two less strict coefficients, we selected association strengths at which the highest percentage of predicted associated pairs carry a known functional association (**S7 Table**). For Yule’s association coefficient Q the cutoff was 0.8, while for Yule’s coefficient of colligation Y it was 0.5. Interestingly, this resulted in us obtaining the same set of 352 putatively contingent gene pairs for both the Q and Y coefficients. We did not set a threshold of significant contingencies for the most stringent phi coefficient of contingency φ, which identified 360 putatively contingent loci pairs, including 351 of the 352 pairs identified by the other two coefficients (above the corresponding threshold). We thus consider the 351 pairs agreed upon by all three metrics as our most reliable set of putatively contingent loci pairs.

When we consider only these 351 most reliable pairs, we still find that 91 out of our 268 (34.0%) convergently mutated loci are involved in at least one putative contingency. This demonstrates that our finding of more than a third of convergently mutated loci being involved in at least in one putative contingency is insensitive to the thresholds applied for our identification of putatively contingent loci pairs.

### Directionality of contingency can be determined for a majority of the most reliable putatively contingent loci pairs

When mutations within one member of a putatively contingent loci pair always occur prior to mutations within the second member, it implies that mutations within the second locus are contingent on mutations within the first one. This logic enabled us to identify the likely directionality of contingency for 294 out of the 351 (83.8%) most reliable putatively contingent loci pairs (see **Materials and Methods, Fig 4A** and **S8 Table**). For an additional 30 pairs, we could not identify the directionality, because mutations within the two loci are always seen together (**S8 Table**). In the remaining 27 putatively contingent loci pairs, directionality could not be assigned because in some instances mutations were first observed in one member of the pair, while in others they were first observed in the second member (**S8 Table**). That for a majority of putative contingencies directionality is consistent and can thus be determined also adds credibility to our predictions.

**Fig 4.**
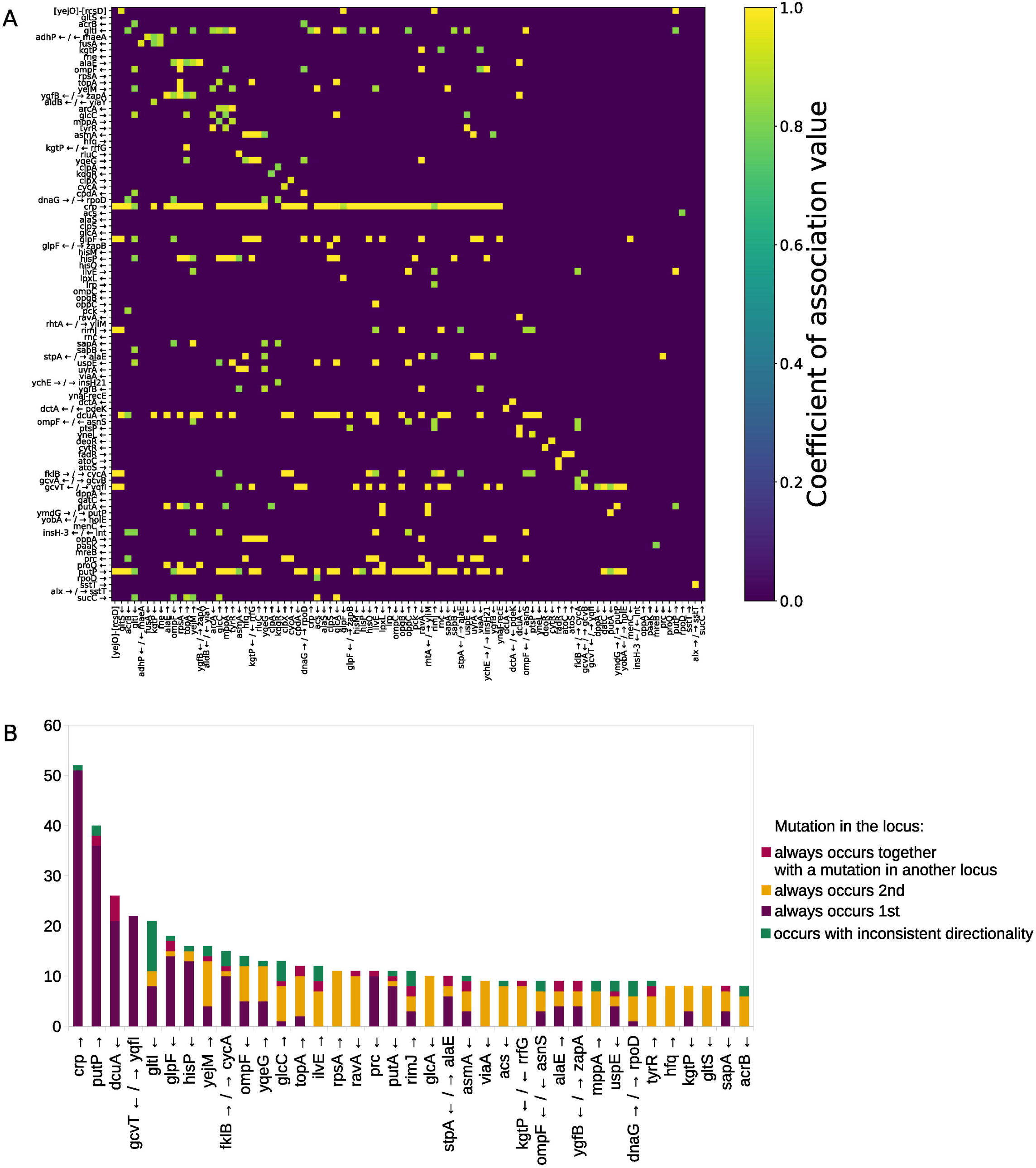
Identified putatively contingent loci pairs. (**A**) Presented are the Yule’s Q coefficient strengths of association of the 351 most reliable putatively contingent loci pairs. These data are represented in a semi-directed adjacency matrix heatmap, in which only in cases where the directionality could not be determined the association is shown as undirected, while for the majority of cases the association is shown as directed. Both rows and columns represent the 91 loci involved in at least one out of the 351 putatively contingent loci pairs. Rows and columns are sorted in the same order. The rows (y axis) represent the loci that were found to be mutated first in each contingency pair, while columns (x axis) represent the loci that were mutated second. (**B**) For each locus that participates in eight or more putatively contingent loci pairs we present a summary of the directionality identified for the putative contingencies in which that locus participates.

### LTSP contingencies with known effect

We identified two putative contingencies for which we already know the biological meaning from our previous studies [28,37]. The first contingency involves mutations within the gene *dnaQ*, which encodes a proofreading component of the DNA polymerase. Such mutations to *dnaQ* occur within two of our populations and occur only on the background of mutator clones. We have previously shown that the combination of a mutation to a mismatch repair gene, with a mutation to *dnaQ* significantly increased mutation rates beyond those of “normal” mutators [28]. The second example involves mutations within the genes *atoC* or *atoS*, that together encode a two component system, which regulates fatty acid metabolism. Such mutations occur exclusively on the background of mutations to the gene encoding FadR - the *E. coli* master regulator of fatty acid metabolism. We have previously shown that this mutation combination allows *E. coli* to metabolize the short chain fatty acid butyrate, which *E. coli* itself produces during the first 24 hours of growth within fresh LB, but which wildtype *E. coli* cannot metabolize [37].

### Mutations to many loci are contingent on previous mutations to the gene encoding the cAMP-activated global transcriptional regulator CRP

We identified some loci that serve as hubs of contingency, meaning that they are involved in many putative contingent loci pairs. When directionality was assigned, it became clear that mutations to the other loci tended to more often be contingent on mutations to these hubs, rather than vice versa (**Fig 4B, S9 Table**). The most striking example of this involves the gene *crp*, which encodes the cAMP-activated global transcriptional regulator CRP and is involved in a total of 52 of the most reliable contingent loci pairs. In 51 of these pairs, mutations to the other locus involved in the pair are putatively contingent on mutations to *crp*, while for the remaining pair the directionality could not be assigned.

We identified 11 putatively contingent loci pairs, involving a transcription factor (TF) and its known target. In 8 of these 11 pairs, mutations within the gene regulated by the TF were contingent on the occurrence of previous mutations within their regulating TF. For seven of these eight pairs the regulating TF was CRP. For six of these seven pairs in which mutations to genes regulated by CRP were contingent on previous mutations to *crp*, the mutations within the regulated loci were never seen in the absence of mutations to *crp*. In the remaining locus mutations are seen in the absence of mutations to *crp* in only one lineage, while they are seen together with mutations to *crp* in six lineages.

Loci that are regulated by CRP, and for which mutations are contingent on previous mutations to *crp* include: (1) the *glpF* gene and its promoter - both mutations in to the *glpF* gene and to its promoter are strongly contingent on previous mutations to *crp. glpF* encodes a glycerol uptake gene, while CRP upregulates *glpF* [41]. Glycerol can be used as an energy-poor carbon source by *E. coli* and the downstream metabolites of glycerol can participate in glycolytic as well as gluconeogenic reactions [42]; (2) *ompF* – is upregulated by CRP and encodes the outer membrane porin F, which is responsible for small molecule diffusion across the membrane [43]. (3) *acs* - encodes an acetyl-coenzyme A synthetase and is upregulated by CRP [44]; (4) *pck* - encodes a phosphoenolpyruvate carboxykinase and is involved in gluconeogenesis. *pck* can be both upregulated or downregulated by CRP [45]. (5) *hfq* - is downregulated by CRP [46] and encodes a key chaperone of RNA folding that is involved in the regulation of interactions between small non-coding RNAs and their targets. As an important chaperone, *hfq* is involved in response to a large variety of conditions and stresses [47,48]; (6) *glcC* – encodes a TF and is upregulated by CRP [49]. GlcC itself regulates the *glcA* gene, which is responsible for the uptake of glycolate [50]. Glycolate can be used by *E. coli* as a carbon source and oxidized to carbon dioxide [51]. Mutations to *glcA* are contingent on previous mutations to *glcC*. This is therefore an example of a chain of contingencies involving regulated genes and their TFs – mutations to *glcA*, which is regulated by GlcC, are contingent on mutations to *glcC*, while mutations to *glcC*, which is regulated by CRP, are contingent on mutations to *crp*.

### Additional putatively contingent loci pairs involving loci with known regulatory interactions

There are several examples of putatively contingent loci pairs that do not involve mutations to *crp*, but nevertheless have known regulatory interactions. Such putatively contingent loci pairs that involve a TF and its target include: (1) the *gcvT*-*gcvA* loci pair - mutations to the gene encoding the GcvA TF are contingent on previous mutations occurring within its regulated gene *gcvT*, which encodes a protein that functions in the glycine cleavage system [52]. For both *gcvA* and *gcvT* mutations occur within their promoter regions, rather than within the genes themselves. (2) *putA* and *putP* – PutA is a TF that downregulates *putP*, which encodes a sodium/proline symporter [53]. No directionality could be assigned to the putative contingency involving these two genes. Mutations were observed within both the *putP* promoter region and the gene itself, in both cases with strong association to mutations in the *putA* gene.

In addition to putative contingencies involving genes and their TFs, we also observed four instances in which mutations to a certain gene were associated with mutations to the same gene’s promoter region: (1) *dctA* and its promoter - Mutations within the *dctA* gene, which encodes a C4-dicarboxylate transport protein, and its promoter are only ever seen together. (2) SstT is a serine/threonine transporter. Mutations in the promoter of *sstT* are contingent on previous mutations to the *sstT* gene itself. (3) mutations to the *putP* promoter region are contingent on previous mutations occurring within the *putP* gene itself. As previously described, *putP* encodes a sodium/proline symporter. (4) mutations to the *cycA* gene, which encodes a D-serine/D-alanine/glycine transporter, only occur on the background of mutations to its promoter. Interestingly, all four examples of putative contingent loci pairs involving a gene and its promoter involve transporter genes (although, as already stated, there is a general enrichment in genes related to transport among the loci that are convergently mutated in our experiments).

Finally, a putatively contingent pair in which both genes co-regulate the same operon was observed. DeoR and CytR co-regulate the *deo* operon, which is involved in nucleoside and deoxy-ribonucleoside catabolism [54]. Interestingly *deoR* and *cytR* are almost always mutated together. Clones that carry mutations to both *deoR* and *cytR* arise early, across all five populations and always belong to the minority of clones that cannot be assigned to any of the lineages, which are all defined by RpoB and RpoC mutations. Clones carrying different mutation combinations to *deoR* and *cytR* can often be seen within a single population (**S2 Table**), suggesting that this mutation combination occurs very frequently. However, they can never establish high frequency lineages, that persist across multiple time points. Combined, this suggests that clones carrying this putatively contingent mutation combination may have a very strong fitness advantage early on under LTSP, but are not able to persist at frequencies above our detection level.

## Discussion

Our results show that *E. coli* adaptation under LTSP continues via highly convergent mutation accumulation at least up to six years into our experiments. We were able to discern the lineage structure of our populations, demonstrating that it is in itself convergent. This means that across populations similar lineages, established by the acquisition of the same lineage defining RNAPC mutations, emerge. Lineages defined by the same RNAPC mutation often behave similarly across populations. The identified lineage structure of our LTSP populations and the high convergence with which mutations are acquired across independently evolving lineages allowed us to identify hundreds of loci pairs for which the acquisition of mutations is statistically significantly associated. Our results demonstrate that contingency is remarkably common, as more than a third of convergently mutated loci are involved in at least one putative contingency.

Our approach has a number of limitations. First, while we can deduce the directionality of 83.8% of the putative contingencies we observed (**S8 Table**), we sometimes cannot assign directionality to our predicted contingencies. A second limitation in our approach is that we cannot tell apart direct contingencies (e.g. mutations in loci A and B are more likely to be adaptive on the background of mutations in locus C) from indirect ones (e.g. adaptive mutations in locus A are more likely to occur on the background of adaptive mutations in locus B, which in turn are more likely to occur on the background of adaptive mutations in locus C). A third limitation of our approach is that it can only identify putative contingent pairs in which both loci involved accumulate mutations convergently, in at least two independently evolving lineages. Instances in which a specific mutation combination occurs in only a single lineage will not be identified by our approach. This means that we would not expect to identify rare contingencies of the type that led to the evolution of citrate utilization in the LTEE [8,11–14]. Finally, as with any statistical analysis false positives are also possible, and thus some of the identified putatively contingent loci pairs may not reflect true contingencies. Indeed, depending on the thresholds and metrics used, the number of contingent pairs identified can vary substantially. This suggests that some of the contingent pairs we identified, for which the association was significant, but not very strong may be less reliable than others. However, the fact that irrespective of the metric and threshold used, the identified putative contingent loci pairs are enriched for known functional associations, suggests that many of these putative contingencies do reflect true biology. Also, irrespective of the metric used, a similarly high percentage of convergently mutated loci (more than one third) were found to be involved in at least one putative contingency relationship, thus demonstrating that our major finding, that contingencies are highly frequent is fairly stable and reliable. Finally, when it comes to the 351 putatively contingent pairs we identified to be the most reliable, for a majority we were able to identify the directionality of the contingency. This means that for these pairs mutations within one member of the pair always occurred prior to mutations in the other. This further adds to the reliability of these putatively contingent loci pairs.

Two non-trivial aspects of the dynamics of adaptation under LTSP enabled us to utilize data extracted from LTSP evolutionary experiments to identify pairs of putatively contingent loci. The first such aspect is the remarkable convergence with which adaptation occurs within our LTSP populations. Adaptation via convergently occurring mutations is a feature of many evolutionary experiments [17–19]. Yet, such convergence is particularly prevalent in our LTSP experiments, where almost 90% of the mutations that occur within non-mutators fall within loci that are mutated across multiple populations and lineages. In addition, we also frequently observed the occurrence of precisely the same mutations across independently evolving lineages and populations. Such specific convergent mutations are less frequently observed in evolutionary experiments [20,55,56]. That convergence is so high in our experiments can stem from strong selection, resulting from intense competition. Under LTSP, only a small fraction of the population is able to replicate, via the recycling of the remains of their ancestors and brethren [27,33]. This likely leads to intense competition between clones. As a result, it seems that the majority of mutations accumulating within nonmutator clones are adaptive (which is also reflected by the very high dN/dS ratios). Strong selection by itself would not necessarily lead to such high convergence, unless the possible pathways to adaptation are greatly limited in number. However, when combined, strong selection and a limited number of possible initial pathways to adaptation can lead to high convergence, of the type we observe.

The second aspect of the dynamics of adaptation under LTSP that contributed to our ability to identify putatively contingent loci pairs was our populations’ lineage structure. We found that lineages are established under LTSP and persist for prolonged periods of time (**Fig 3B**), while continuously accumulating mutations (**S2 Fig**). Such a lineage dynamic indicates that the cells that constitute the majority of the replicating sub-population continuously belong to the same lineages. In other words, it demonstrates that most cells that are able to replicate at a given time point are decedents of those that were able to replicate at previous time points. A contrary example to this general trend, which demonstrates that such a dynamic is far from trivial, involves clones carrying a putatively contingent mutation combination in the *cytR* and *deoR* genes. We observed such clones early on in our experiment, across all five of our populations. Unlike the great majority of clones sampled during our experiment, these clones did not carry an RNAPC lineage-determining mutation. Clones with a *cytR* + *deoR* mutation combination did not establish lineages of their own. Instead, we observed clones carrying specific mutation combinations within these loci rising to relatively high frequencies at a certain early time point, only to never be seen again at a later time point. Clones carrying a mutation combination within *cytR* and *deoR* that were sampled from the same population at later time points included an entirely different mutation combination within these genes. Thus, it can be said that for the clones carrying a *cytR* + *deoR* mutation combination the cells that replicated and rose to high frequencies at a given time point, were not decedents of those that replicated and rose to high frequencies during a previous time point.

Contingencies are widely thought to lead to increasing diversification in adaptive pathways [3,15,16]. After all, if different lineages acquire different adaptations early on, their adaptive trajectory may continue to diversify due to different adaptations being contingent on these initial adaptations. Given that we observe frequent putative contingencies, we may thus expect to see in our populations a pattern of reducing convergence with time. Fitting with this we indeed observe that while still highly convergent, non-mutator mutations that accumulate at later time points in our experiment appear to be less convergent than those occurring early on (**S1C Fig**). While such a pattern could also be explained by weaker selection to adapt at later time points, there is no a-priori reason to think that in our experiment selection would be weaker at later time points. Indeed, as time progresses, our populations’ carrying capacity reduces (meaning that the number of cells allowed to persist diminishes, **Fig 2A**), which may lead to even stronger competition between clones. Furthermore, across time points dN/dS of non-mutator mutations remains significantly higher than 1 (**S10 Table**), suggesting continuous adaptation under positive selection. While the convergence of adaptation within our populations does seem to reduce somewhat with time, adaptation is still highly convergent even six years into our experiments. Even in the last time point sampled, 34.9% of mutations occurring within non-mutators fall within loci that are mutated across all five populations and 84% fall within loci mutated in at least two.

While historical contingencies are thought to contribute to diversification of adaptive trajectories, very frequent contingencies can also have the somewhat opposite effect of increasing population-level parallel adaptation. After all, if initially the same adaptations are often sampled across populations and strongly influence the identity of future adaptations, this may contribute to similar adaptive trajectories arising across populations. Overall, the ability to observe such convergent adaptive trajectories will depend on the number of clones one samples from each population and timepoint. Specific clones may adapt very differently from other clones present within their populations and variation in adaptations acquired by individual clones may greatly increase with time, both by chance and due to contingencies. Thus, if researchers sequence only one or two clones per population and time point sampled, as is often done following evolutionary experiments, they will likely observe large variation between populations. At the same time, by sequencing ∼10 clones per population and time point sampled, we were able to observe that even though individual clones may adapt via highly divergent trajectories, at the population level one can observe very high levels of parallel adaptation. It is however important to note that by sequencing ∼10 clones out of many millions, we also could identify only the most frequent genotypes present within our populations. Convergent adaptations present at slightly lower frequencies within some or all populations may have thus been missed entirely, or not have been assigned as occurring convergently across multiple populations or lineages. It is thus possible that the extremely high levels of convergent adaptation we observed constitute an under-estimate of the true extent of convergent adaptation under LTSP.

To conclude, we demonstrate that LTSP populations adapt in an extremely parallel manner, with a majority of mutations occurring within non-mutator clones falling within loci that are mutated across multiple populations. One feature of the parallelism with which LTSP populations adapt is the establishment and persistence of lineages that tend to behave with similar dynamics across populations. Combined, the convergence with which adaptive mutations occur and the lineage structure of our populations enables us to identify hundreds of instances of putative historical contingencies and reveal that such contingencies are very common, affecting more than a third of convergently mutated loci.

## Materials and Methods

### Sequencing of LTSP clones and mutation calling

One milliliter of each of the five LTSP populations, established in July 2015, was sampled at days 0, 1, 2, 3, 4, 5, 7, 8, 11, 22, 32, 42, 53, 64, 94, 127, 154, 184, 241, 308, 374, 436, 557, 730, 1095, 1178, 1460, 1612, 1816 and 2191. Sample dilutions were used to estimate viability by quantification of colony-forming units (CFUs) with the help of robotic plate reader. Samples were preserved in 50% at -80°C for further analysis. Dilutions of samples from days 11, 22, 32, 42, 64, 127, 374, 730, 1095, 1460, 1816 and 2191 were plated and cultured overnight. ∼10 colonies from each cultured dilutions were inoculated in 4 ml of media and were subsequently cultured until the optical density of 1 was reached. DNA was extracted with the Macherey-Nagel NucleoSpin 96 Tissue Kit from pelleted cells, resulting from the centrifugation of one milliliter of each culture at 10 000 g for 5 minutes. The remainder of each culture was frozen in 50% at -80°C. Libraries were prepared according to the protocol described in [57]. 150 bp paired end sequencing was carried out on an Illumina HiSeq 2500 machine at Admera Health. Clones from all five populations from days 1460, 1816 and 2191 as well as clones from time points 22, 32 and 42 from populations 3 and 5 were sequenced during the current study, while clones from all other time points and the ancestral clones were sequenced as part of our previous studies **(S1 Table)** [27,28]. Obtained reads were aligned to the *E. coli* K12 MG1655 reference genome (accession NC_000913) and mutations were called, using Breseq [58], which enabled us to identify SNPs as well as small and large indels (**S2 Table)**.

Raw short read sequencing data obtained for this study were deposited to the Sequence Read Archive (SRA) under accession number PRJNA1049885. Short read data obtained as part of our previous studies can be found under accessions BioProject ID: PRJNA380864 and BioProject ID: PRJNA674613.

### dN/dS calculations

Numbers of synonymous and non-synonymous sites of all protein coding sequences of *E. coli* were calculated as in [27]. Briefly, The protein-coding gene sequences of *E. coli* K12 MG1655 were obtained from the NCBI database. For each DNA position of each protein coding gene, we calculated the likelihood that a mutation within that site would be synonymous or non-synonymous. These likelihoods were then summed up across all protein coding sequences. The dN/dS ratio was then calculated for non-mutator clones as dN/dS =

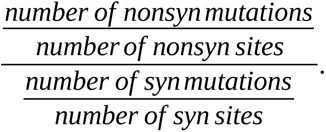

### Building phylogenies within each LTSP population

To characterize the lineage structure of each LTSP population, we created a database of the mutations found within each of its sequenced clones. Only mutations that rose to a frequency of 30% and higher at any time point throughout the experiment were included in the databases. Using these data, the genetic distances between clones were estimated using the Levenshtein distance metric, while perceiving mutations as single characters and lists of mutations as strings (the Levenshtein distance is a measure of the difference between two strings that is estimated as the minimal number of character replacements needed in order to convert one string to the another [59]). The obtained distances were utilized to determine the “parent” for each clone and the resulting ancestor-descendant pairs were used to build phylogenies for each of the five populations and create Muller plots (**Fig 3B**) with the help of the R package MullerPlot [36].

### Gene Ontology Enrichment Analysis

To test whether loci mutated across two or more lineages are enriched for particular biological processes, we used the Gene Ontology’s tool “GO Enrichment Analysis” [60–62]. The list of the genes used in the analyses can be found in **S4 Table**. Statistically significant enrichment was deduced when the FDR corrected P value was smaller than 0.05.

### Screening for putatively contingent loci pairs

We focused on 268 loci that were mutated in at least two independent lineages or populations, excluding *rpoB* and *rpoC*. From this list of loci we generated a list of pairs of loci that were mutated together, within a non-mutator clone in at least two lineages or populations. For each such pair of loci, we then generated contingency tables by counting in how many lineages did we see clones that carried mutations to both loci, clones that carried mutations to neither locus, and clones that carried mutations in only one or only the other of the two loci. It should be noted that the same lineage can be counted towards different categories simultaneously, meaning that for example we can see within a specific lineage clones in which both loci are mutated, and others in which only one of the two is mutated. In such a case, for this pair of loci, that lineage will be counted under both categories. In order to account for contingencies that occur within the rare clones that could not be assigned to any lineage, we assigned such clones to “pseudo-lineages” (one for each population in which such clones existed). In total, after removing the three mutator lineages, there remained a total of 20 lineages and pseudo lineages.

Using the obtained contingency tables, we calculated three metrices of association: Yule’s association coefficient Q [38], Yule’s coefficient of colligation Y and Yule’s phi coefficient of contingency φ [39]. To ascertain statistical significance, we performed a one-sided t-test with α=0.05 followed up with FDR multiple testing correction (**S5 Table**).

### Functional association analyses

We used the STRING database [40] to quantify the fraction of putatively contingent loci pairs that are already known to be functionally associated. To examine whether this number was higher than expected by chance, we performed 1000 randomizations where we selected at random loci pairs from the list of 268 convergently mutated loci. For each randomization we could then quantify what fraction of randomly selected loci pairs were functionally associated according to the STRING database. These numbers were then used to generate a random distribution and calculate a P-value for the observed numbers of putatively contingent loci pairs that were also functionally associated. 1000 randomizations were carried out separately for each of the three metrices of association used, and in each instance the number of loci pairs selected per randomization matched the number of putatively contingent loci pairs identified using that metric.

### Estimation of contingency directionality

In order to estimate the directionality of the contingency for each of the most reliable 351 putatively contingent loci pairs, we considered only the lineages in which mutations to both members of the pair were found within single clones. For such clones the phylogenies were built by tracking the chain of ancestor-descendant pairs of clones with the smallest Levenshtein distance, while representing mutations as single characters and all mutations in the clones as strings. The obtained data was used to examine whether the predicted contingent mutations always occurred together in the same clones of the lineage or whether one of the mutations could occur separately from another. In cases where the mutations did not always occur together, the directionality was determined by identifying which of the pair of mutations was found earlier in an ancestor of the clone that eventually carried both putatively contingent mutations. The resulting data was then combined across lineages to determine the ultimate trend (e.g. that across lineages one of the mutations was always seen prior to the other (**S8 Table**).

## Data availability statement

Raw sequencing reads were deposited to the sequence read archive (SRA) BioProject ID: PRJNA1049885.

## Supporting information captions

**S1 Table. Obtained sequence coverage**

**S2 Table. Full list of called mutations**

**S3 Table. Loci mutated convergently in two or more populations**

**S4 Table. Loci mutated convergently in two or more lineages**

**S5 Table. Strength of association of putatively contingent loci pairs**

**S6 Table. Summary of 1000 simulations of contingent loci pair networks**

**S7 Table. Characteristics of various association coefficient value cut-offs**

**S8 Table. Directionality of putatively contingent loci pairs**

**S9 Table. Contingency directionality summary for each locus involved in at least one putatively contingent loci pair**

**S10 Table. dN/dS of non-mutator mutations across time points**

**S1 Fig. Highly convergent pattern of mutation acquisition.**

The charts show the fractions of mutations occurring in non-mutators (A) across populations; (B) across lineages; (C) across population as a function of time. In the chart C the mutations are counted towards the category at each time point if it has been observed in n populations at and up to the time point.

**S2 Fig. Mutation accumulation per lineage in three LTSP populations of E. coli that were not overtaken by mutators.**

Data from the RpoB E1272G mutator lineages are not presented. Dots represent mean values across clones within each lineage with error bars representing standard deviations around these means.

