## Supplementary figures and images for "*Escherichia coli* adaptation under prolonged resource exhaustion is characterized by extreme parallelism and frequent historical contingency"

### S1 Fig

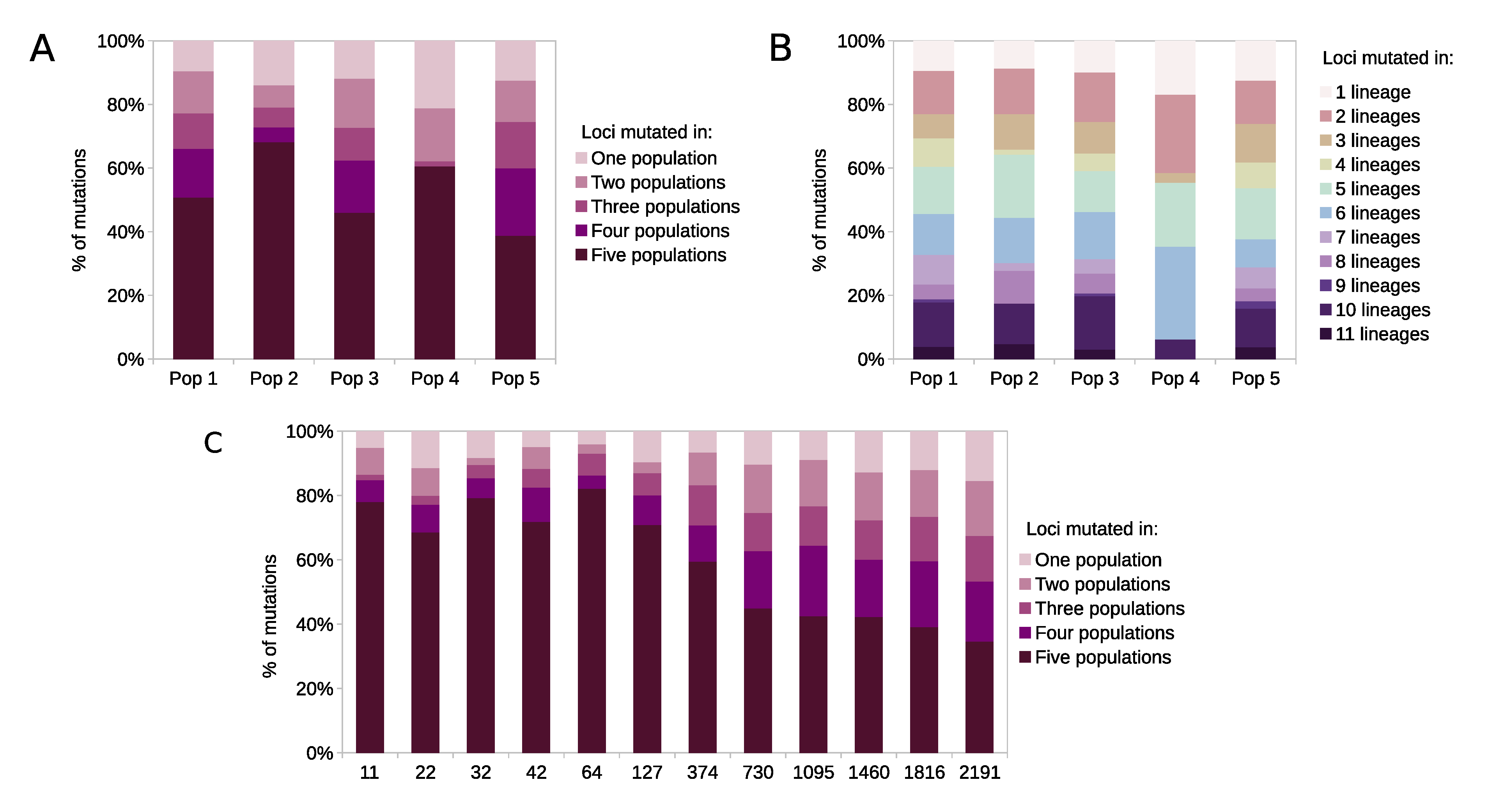

### S2 Fig

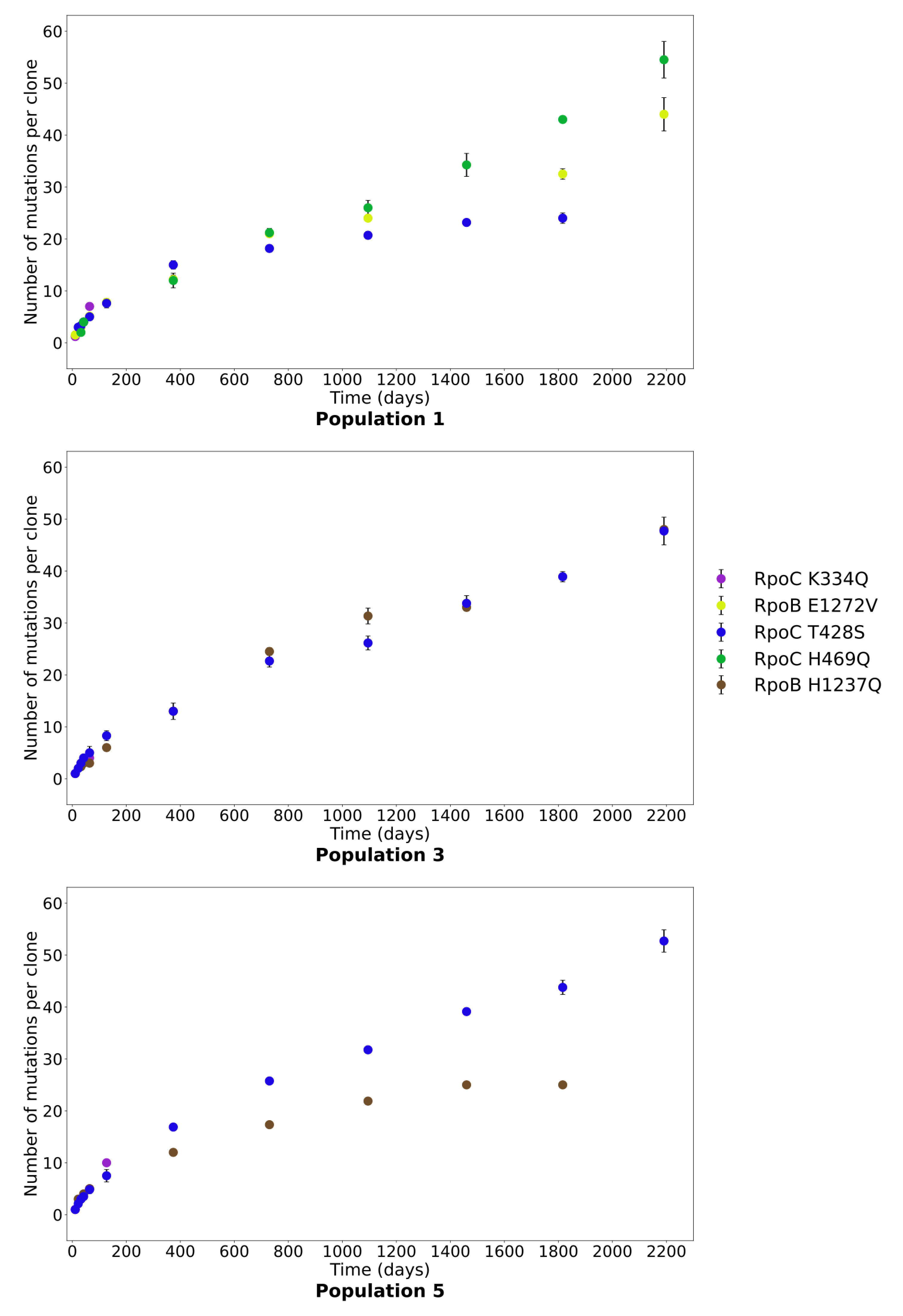
